# Unmixing Biological Fluorescence Image Data with Sparse and Low-Rank Poisson Regression

**DOI:** 10.1101/2023.01.06.523044

**Authors:** Ruogu Wang, Alex A. Lemus, Colin M. Henneberry, Yiming Ying, Yunlong Feng, Alex M. Valm

## Abstract

Multispectral biological fluorescence microscopy has enabled the identification of multiple targets in complex samples. The accuracy in the unmixing result degrades (1) as the number of fluorophores used in any experiment increases and (2) as the signal-to-noise ratio in the recorded images decreases. Further, the availability of prior knowledge regarding the expected spatial distributions of fluorophores in images of labeled cells provides an opportunity to improve the accuracy of fluorophore identification and abundance. We propose a regularized sparse and low-rank Poisson unmixing approach (SL-PRU) to deconvolve spectral images labeled with highly overlapping fluorophores which are recorded in low signal-to-noise regimes. Firstly, SL-PRU implements multi-penalty terms when pursuing sparseness and spatial correlation of the resulting abundances in small neighborhoods simultaneously. Secondly, SL-PRU makes use of Poisson regression for unmixing instead of least squares regression to better estimate photon abundance. Thirdly, we propose a method to tune the SL-PRU parameters involved in the unmixing procedure in the absence of knowledge of the ground truth abundance information in a recorded image. By validating on simulated and real-world images, we show that our proposed method leads to improved accuracy in unmixing fluorophores with highly overlapping spectra.

## 1 Introduction and motivation

Many biological systems are composed of multiple interacting subcomponents, any of which may be labeled with fluorescent reporters to map their spatial location within cells and tissues. While some progress has been achieved in developing fluorescent dyes with narrow emission spectra, e.g., semiconductor nanocrystals or quantum dots, most widely used organic fluorophores, including fluorescent proteins, have broad excitation and emission spectra [Zimmermann, 2005, Barroso, 2011]. The inherent wide emission spectra of different fluorophores used in a single experiment lead to unavoidable overlap in spectral emission profiles and cross-talk between the recorded channels when samples are imaged with conventional bandpass filters. To overcome this limitation, fluorescence spectral imaging instrumentation and image analysis tools have been developed and applied to biological imaging. Spectral imaging microscopes collect fluorescence intensity information at every pixel in an image to construct a 3-dimensional data cube with spatial and spectral information from the sample as shown in Figure 1.

**Figure 1:**
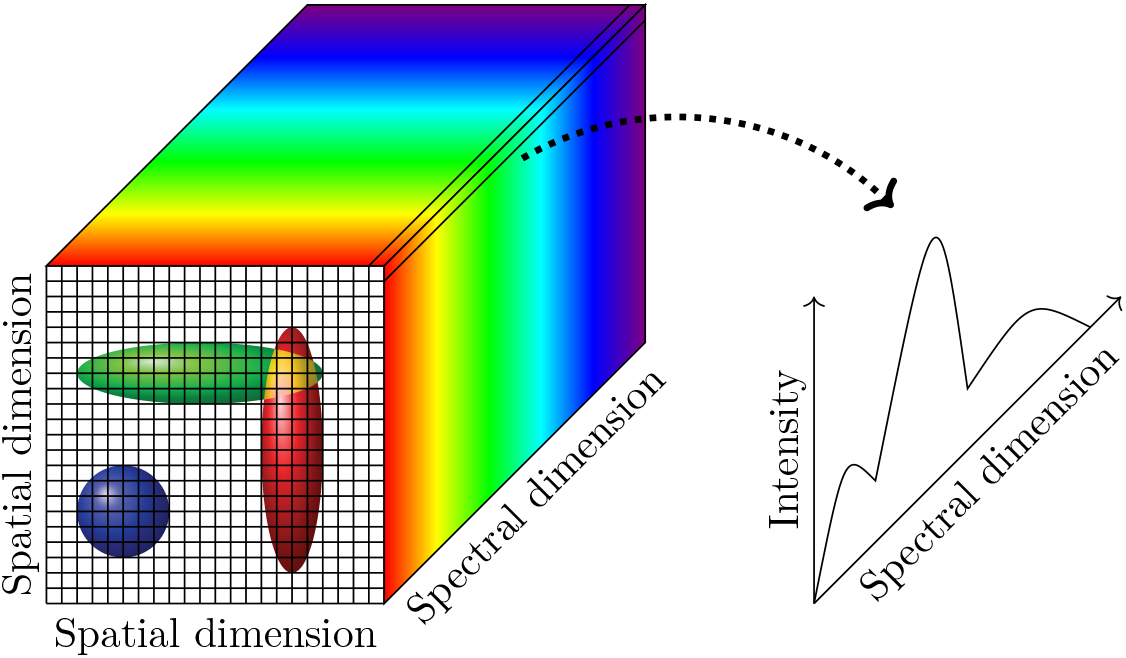
A hyperspectral data cube and spectral intensity information at each pixel with a generalized biological image, in which most foreground pixels record fluorescent signal from only one cell but some pixels record overlapped signals from two or more cells.

Among existing biological spectral imaging analysis methods, spectral unmixing, which aims at extracting the spectral signature of each fluorophore from recorded images and gaining knowledge of each fluorophore’s abundance in every pixel, has been widely utilized. In particular, linear unmixing approaches, especially utilizing the least squares framework, have been widely applied since these approaches make no underlying assumptions about the image data except that the signal recorded in the same pixel from multiple fluorophores adds linearly, i.e., no a priori knowledge about the sample is required. Linear unmixing separates each pixel linearly into the spectral signatures of the contributing fluorophores, called *endmembers*, and their contributions, called *abundances.* Due to physical considerations, both endmembers and abundances should satisfy the nonnegativity constraint. Given a spectral image matrix 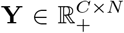 with *C* channels and *N* pixels, denoting 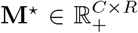 as the endmember matrix of the associated *R* fluorophores and 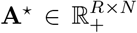 the corresponding abundance matrix, such a linear relationship can be expressed as

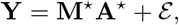

where 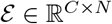 denotes an unknown noise matrix.

Least squares (linear) unmixing procedures are implemented by decomposing a spectral image matrix into an endmember matrix and an abundance matrix simultaneously while minimizing the data fidelity error measured by the squared error criterion, e.g., the sum of the squared residuals. Mathematically, it can be formulated as follows

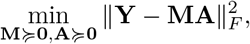

where ||·||_*F*_ denotes the Frobenius norm of a matrix, and 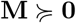 and 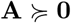 denote element-wise nonnegativity of the endmember matrix **M** and the abundance matrix **A**. In biological spectral imaging, it is frequently the case that reference spectral images that consist of only one fluorophore— and so only one endmember—in each image are available, which is also the scenario considered in this study. In such a case, least squares unmixing procedures can be carried out in a two-step way: first, each reference spectral image matrix is decomposed into the outer product of two vectors, i.e., the endmember and its abundances; then, with the extracted endmember information from the first step, the abundance matrix is estimated for the mixed image. Such a two-step approach is also considered in this study.

While least squares unmixing has been extensively studied and widely used in the spectral unmixing literature, its limitations are also well-recognized in the community. For instance, due to the involvement of matrix multiplication, least squares unmixing may not admit a unique solution, which leads to imprecise estimates of the abundances. In the literature, this problem is frequently addressed by imposing a certain penalty on *A*, which leads to

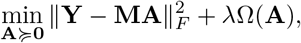

where λ > 0 is a tuning parameter, Ω(**A**) denotes the penalty imposed on **A**, and **M** is assumed to be known from reference images. The penalty term λΩ(**A**) controls the complexity of the matrix space within which **A** is searched and thus may lead to a unique solution. Another disadvantage of the least squares unmixing approach lies in the use of the least squares error criterion, which originates from the maximum likelihood estimation when assuming Gaussian noise. In fact, the presence of purely Gaussian noise is rarely the case in biological spectral images. This is because the spectrum at each pixel of a biological fluorescence image corresponds to the photon counts recorded at every spectral channel and the uncertainties in these measurements can be better approximated by Poisson distributions rather than normal distributions (the so-called, photon shot noise); see e.g., Coates [1972]. Following this observation, Poisson regression approaches to spectral unmixing have been proposed in the literature; see e.g., Neher and Neher [2004], Zimmermann [2005]. To date, existing applications of Poisson regression approaches to biological spectral unmixing comprise an active area of research, and the strengths and weaknesses of such approaches, when being compared with well-established least squares approaches, are yet to be fully elucidated.

Least squares approaches to spectral unmixing are implemented in a pixel-wise fashion, which requires no assumption regarding fluorophore distribution in the image. In biological spectral images, neighboring pixels are frequently similar in their fluorophore identities and abundances. This spatial correlation can be reflected as linear dependence between the corresponding abundances. As a result, the rank of the matrix formed by these abundance vectors may be limited, especially if we consider neighboring pixels in a small region. In addition, for a real biological spectral image, it is frequently the case that the recorded signal from any given pixel may only comprise one or a small number of endmembers, though, for the whole image, the number of involved endmembers may be far larger. However, least squares unmixing tends to treat all pixels and endmembers equally without taking such sparseness information into account, which may lead to imprecise abundance estimation when the number of contributing endmembers in any pixel is much smaller than that of the candidate pool across the entire image.

Here we address these gaps in knowledge and further take advantage of the prior information available regarding spectral images of fluorescently labeled cells by exploring a Poisson regression approach and by simultaneously seeking endmember-wise sparseness and low-rankness when estimating the abundances in a localized pattern. We propose in this paper a regularized sparse and low-rank Poisson regression approach (SL-PRU) to accurately estimate abundances in multiplex labeled images of cells. The proposed approach takes into account the non-Gaussian nature of the noise in the data and neighboring information, which are physically meaningful considerations in biological spectral imaging. To implement the low-rankness assumption of neighboring pixels, we make use of a sliding window technique, which allows us to consider the spectral signatures of adjacent pixels lying in the window. To make SL-PRU computationally tractable and to pursue further robustness, we consider convex relaxations of the penalty terms on the sparsity and the rank of the abundance matrix.

It is noticed from our empirical studies that unregularized Poisson regression may lead to satisfying endmember extraction results. We thus suspend the penalty terms when implementing SL-PRU to extract endmembers in the first step from images with known fluorophore identities. In the second step, where SL-PRU is applied to unmix real biological images, we propose a constructive approach for tuning the parameters involved in the estimation without resorting to the unknown abundance matrix. We validate the proposed method on simulated spectral images and on real images of a microbial biofilm. The experimental results show that our proposed method can outperform existing approaches in biological imaging both quantitatively and qualitatively.

## 2 Materials and methods

### 2.1 Sample preparation

E. coli K12 (ATCC 10798) cells were grown to the mid-log phase in Luria-Bertani LB Broth (Difco Laboratories, Inc.). E. coli cultures and dental plaque smears were fixed in 2% paraformaldehyde (EMS Diasum) for 1.5 hours at room temperature, then stored in 50% ethanol for 24 hours before FISH labeling. E. coli cells were labeled with the general bacteria probe, EUB338 (GCTGCCTCC-CGTAGGAGT) conjugated to a fluorescent dye at the 5’ end (Thermofisher). Plaque smear samples were obtained through self-flossing from healthy volunteers after giving informed consent. The use of human subjects for this study was approved by the University at Albany Institutional Review Board (IRB). Plaque samples were labeled with previously validated taxon-specific FISH probes (See Supplementary Table 1) and acquired as multi-plane z-stack images.

### 2.2 Imaging and pre-processing

Images were acquired on Zeiss LSM 710 or LSM 880 confocal microscopes with 32 anode spectral detectors. Images were acquired with 488, 561, and 633 nm laser excitation and collected on the 32-anode spectral detector with 9.8 nm width spectral resolution in each channel. E. coli images were acquired as a single plane with a 63x 1.4 NA objective. Plaque smear images and reference E. coli images were acquired as multi-plane Z-stack images with a 20x 0.8 NA objective.

### 2.3 SL-PRU: The proposed unmixing approach

Since the signal from multiple fluorophores adds linearly in a pixel, we can express the true photon counts as the sum of the endmembers weighted by their abundances. Considering that the recorded photon counts follow the Poisson distribution, a biological spectral image 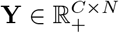 with *C* channels and *N* pixels can be written as [Neher et al., 2009, Novo et al., 2013, Xu et al., 2020]

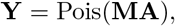

where 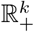 denotes a nonnegative orthant in a *k*-dimensional Euclidean space, Pois(·) is element-wise Poisson probability distribution, 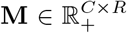 represents the endmember matrix which consists of reference spectra of *R* fluorophores used for labeling, and 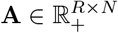 denotes the abundance matrix.

To extract the endmember 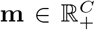 from a reference image 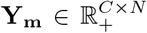, we maximize the likelihood of observing 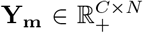 given **m** and the corresponding abundances 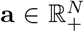, which leads to the following Poisson Nonnegative Matrix Factorization (PNMF):

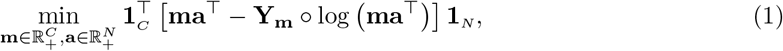

where **1**_*C*_ and **1**_*N*_ are vectors of length C and N whose entries are all 1 and o denotes element-wise multiplication.

With endmembers extracted through reference images via (1), the abundance estimation of a spectral image that shares the same morphologies can be viewed as a nonnegative Poisson regression problem. As both endmember-wise sparsity and spatial correlation properties rely on the homogeneity of a small area, we utilize a sliding 3 × 3 window that contains the spectra of the target pixel and spectra of its surrounding pixels for unmixing as shown in Figure 2 [Giampouras et al., 2016]. In each window, we assume the contribution of only a few fluorophores which can be reflected as a limited number of endmembers with nonzero abundances. In the literature, the sparsity constraint is usually carried out via an *ℓ*_1_ norm regularization that reduces the number of nonzero entries [Iordache et al., 2011, Rossetti et al., 2020]. To impose endmember-wise sparsity on the abundance matrix **A** in a window, we apply the constraint among the rows of **A** which turns out to be the *ℓ*_2,1_ norm 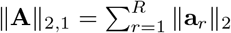 where **a**_*r*_ denotes the *r*-th row of **A** and || · ||_2_ denotes the *ℓ*_2_ norm [Iordache et al., 2013]. The other property, spatial similarity of neighboring pixels, has also been widely exploited for hyperspectral unmixing [Iordache et al., 2012, Giampouras et al., 2016, Zhang et al., 2018]. In this work, the spatial correlation is incorporated by imposing the low-rankness constraint on **A**. As a convex surrogate of the matrix rank, the nuclear norm ||**A**||_*_ defined as the sum of its singular values, 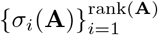, is used.

**Figure 2:**
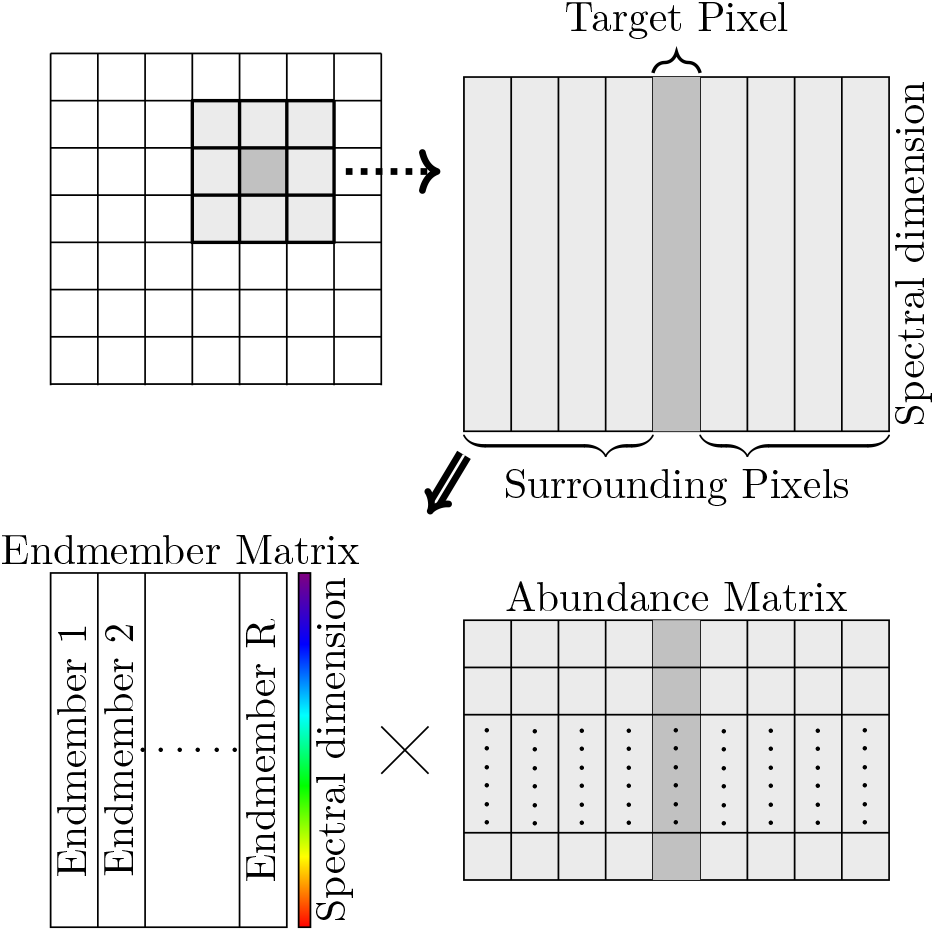
A3 × 3 window unmixed into the product of its endmember and abundance matrices

With a slight abuse of notation, we denote in the following 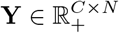 as a spectral window. The abundance matrix **A** can be estimated through the following sparse low-rank Poisson regression:

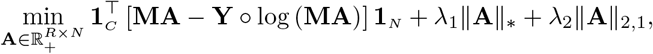

where λ_1_ and λ_2_ are nonnegative parameters that balance the fidelity term, the penalty term on sparsity, and the penalty term on low-rankness.

As convex relaxations of the norm, i.e., the number of nonzero entries of a vector, and the rank function of a matrix, the *ℓ*_1_ norm and the nuclear norm suffer from the influence of the magnitude [Candes et al., 2008, Lu et al., 2014]. We thus use weighted formulations of nuclear norm and *ℓ*_2,1_ norm to democratically penalize the nonzero entries:

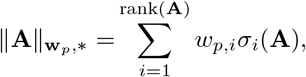

and

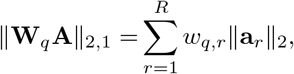

where

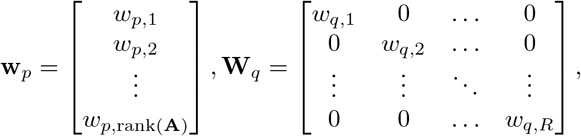

and 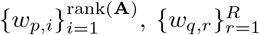 are nonnegative weighting coefficients that will be determined later. As a result, we arrive at the following variant of the above regularized sparse and low-rank Poisson regression method:

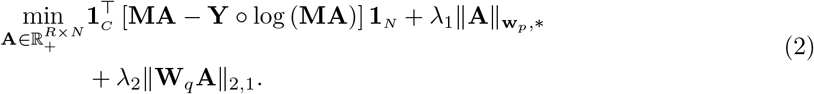

The proposed algorithms for solving PNMF (1) and SL-PRU (2) and their pseudo codes are provided in Supplementary Material.

## 3 Experiments and results

The performance of our proposed approach, SL-PRU, is compared with the commonly used linear unmixing methods: nonnegative least squares approach (NLS) [Zimmermann, Timo, 2005], sparse NLS (S-NLS) [Bioucas-Dias and Figueiredo, 2010, Rossetti et al., 2020], and sparse and low-rank NLS (SL-NLS) [Giampouras et al., 2016].

We carry out three different sets of experiments in which different types of spectral image data are used. In the first set of experiments, we unmix reference images of E. coli cells, in which every cell in the image contains a single endmember of knonw identity. Therefore, the performance of the different approaches in unmixing this data set can be compared by evaluating the proportion of the correct endmember assigned to each foreground pixel in the images while assuming all endmembers are also involved in each image. In our second set of experiments, we consider simulated spectral images, which are generated by using simulated abundance matrices and uncorrelated and correlated endmembers and by adding Poisson noise with different signal-to-noise ratios (SNRs). The root mean square errors (RMSEs) of the unmixing solutions from the different methods are evaluated for comparison. As a third set of experiments, we also evaluate the effectiveness of our proposed approach on real biological spectral images of labeled microbial biofilms and compare the results with NLS.

**Table 1:**
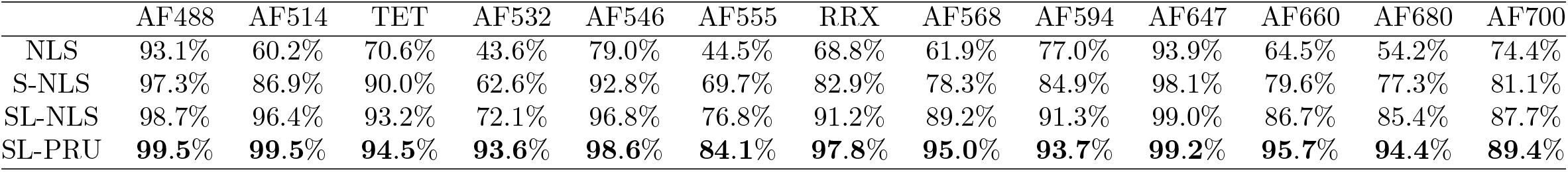
Optimal average proportions of endmembers estimated by NLS, NLS with sparsity constraint (S-NLS), NLS with sparsity and low-rank constraints (SL-NLS), and Poisson regression with sparsity and low-rank constraints (SL-PRU).

For all three experiments, the endmember spectra used for abundance estimation are extracted through PNMF (1) from known reference images. The tuning parameters λ_1_ and λ_2_ in (2) are chosen from the set {0,10^-3^,10^-2^, 10^-1^,1,10}, and *μ* in Algorithm for solving SL-PRU (See Algorithm 2 in Supplementary Material) is fixed to 0.01.

### 3.1 Reference images: endmember extraction and unmixing

We first extract endmembers using PNMF (1) on thirteen reference images of labeled E. coli cells and apply SL-PRU to unmix them to estimate abundances. The standardized fluorometer measured spectra of thirteen endmembers against wavelength and the standardized endmember matrices extracted by the arithmetic mean method [Rossetti et al., 2020] and PNMF are displayed in Supplementary Figure 1 for comparison. See Supplementary Table 2 for the full names of all fluorophores and their abbreviations used here.

Given an estimated abundance vector **a** = (*a*_1_, *a*_2_, · · ·, *a_R_*)^*T*^, where *a_r_*, *r* =1, · · ·, *R*, denotes the estimated abundance of the *r*-th endmember of a pixel from a reference image being unmixed. Then, the proportion of the involved endmember in this pixel is defined as

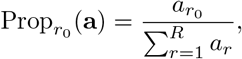

where *α*_*r*_0__ denotes the abundance of this endmember in the pixel. For a reference image, the average proportion (of the involved endmember) is defined as the average of such proportions across all pixels in this reference image. Note that for each reference image, only one endmember is involved. Therefore, the closer the average proportion is to 1, the better the estimated abundance vector.

With each tuning parameter for S-NLS or each pair of tuning parameters for SL-NLS or SL-PRU, we can obtain thirteen average proportions for all thirteen reference images. We choose the minimum of the thirteen average proportions for comparisons, since this represents the most conservative measure of unmixing accuracy. The tuning parameter values that reach the highest minimum of the thirteen average proportions obtained through each method are regarded as the optimal values. The average proportions of thirteen reference images obtained through each method: NLS, S-NLS, SL-NLS, and our proposed model SL-PRU with the optimal parameters are reported in Table 3.1. As shown in Table 3.1, our proposed method provides the highest average proportions among all the methods for all thirteen reference images.

### 3.2 Unmixing simulated spectral images

Using the endmembers extracted from the same reference images as in the first set of experiments above, we simulate spectral images of size 3 × 3 pixels with two uncorrelated endmembers AF514 and RRX and two correlated ones, AF555 and RRX, respectively. The root mean square error criterion is used to compare the performance of different methods, where

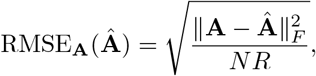

where **A** is the simulated abundance matrix that serves as the underlying truth and ***Â*** the estimated abundance matrix.

#### 3.2.1 Simulation 1: Uncorrelated endmembers

The abundances of AF514 and RRX are generated from the uniform distribution *U* [0,1] and the abundances of other endmembers are set to zero. Then the simulated spectral image is the multiplication of the generated abundance matrix and the endmember matrix. Poisson noise with nine SNRs ranging from 2 to 10 is added to the simulated data, and 1000 realizations of such simulated images are generated for each SNR.

With the optimal parameter(s) being selected for the cases with different SNRs, the averaged RMSEs in 1, 000 repetitions of all the unmixing methods under comparison are plotted in Figure 3. In particular, when SNR is set to 5, the optimal average RMSE is obtained from SL-PRU with λ_1_ =0.1 and λ_2_ = 1. Moreover, the average of the estimated abundance matrices for this scenario is plotted in the top row of Figure 5.

**Figure 3:**
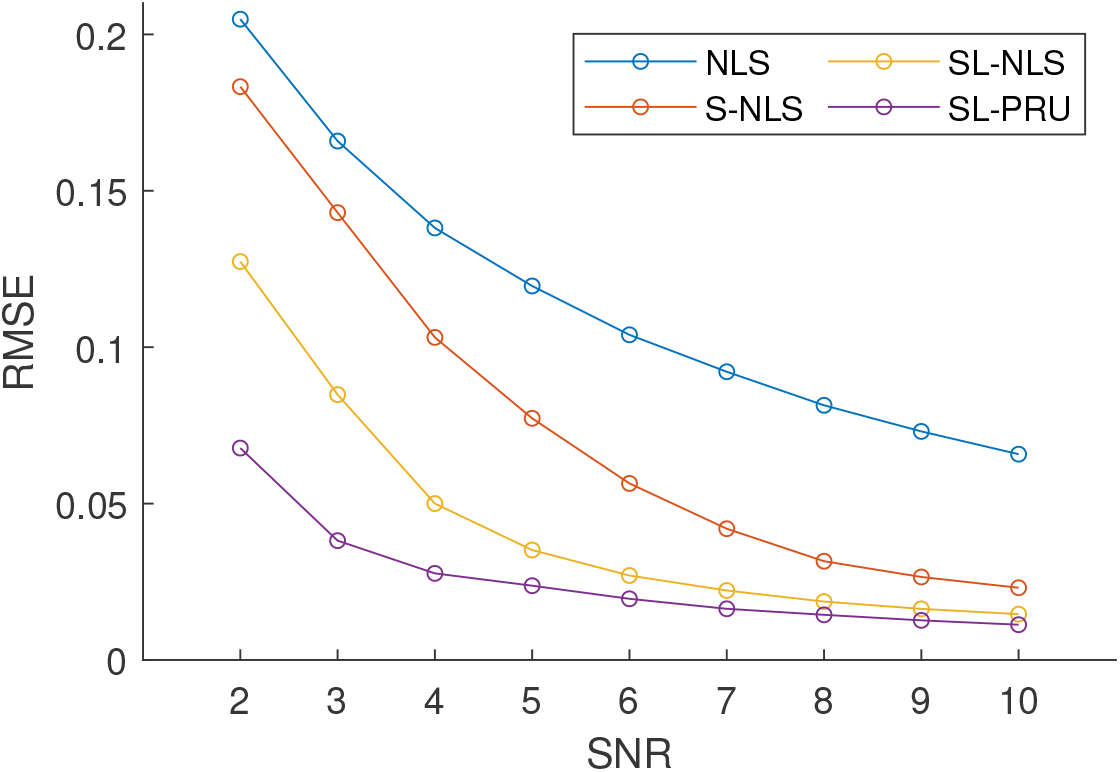
Averages of RMSEs of the abundances estimated by each of the unmixing methods that we considered from simulated images that contain colocalized AF514 and RRX with Poisson noise and SNR of 2 to 10.

**Figure 4:**
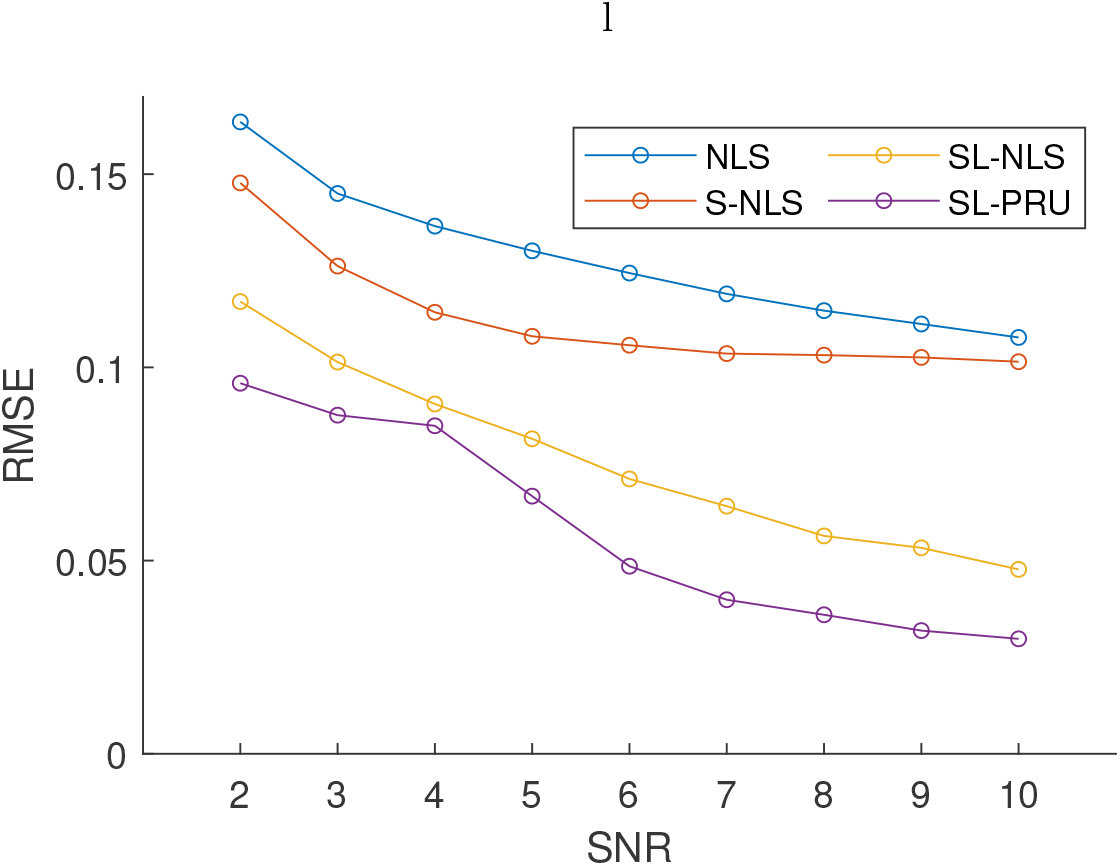
Averages of RMSEs of the abundances estimated by each of the unmixing methods that we considered from simulated images that contain colocalized AF555 and RRX with Poisson noise and SNR of 2 to 10.

**Figure 5:**
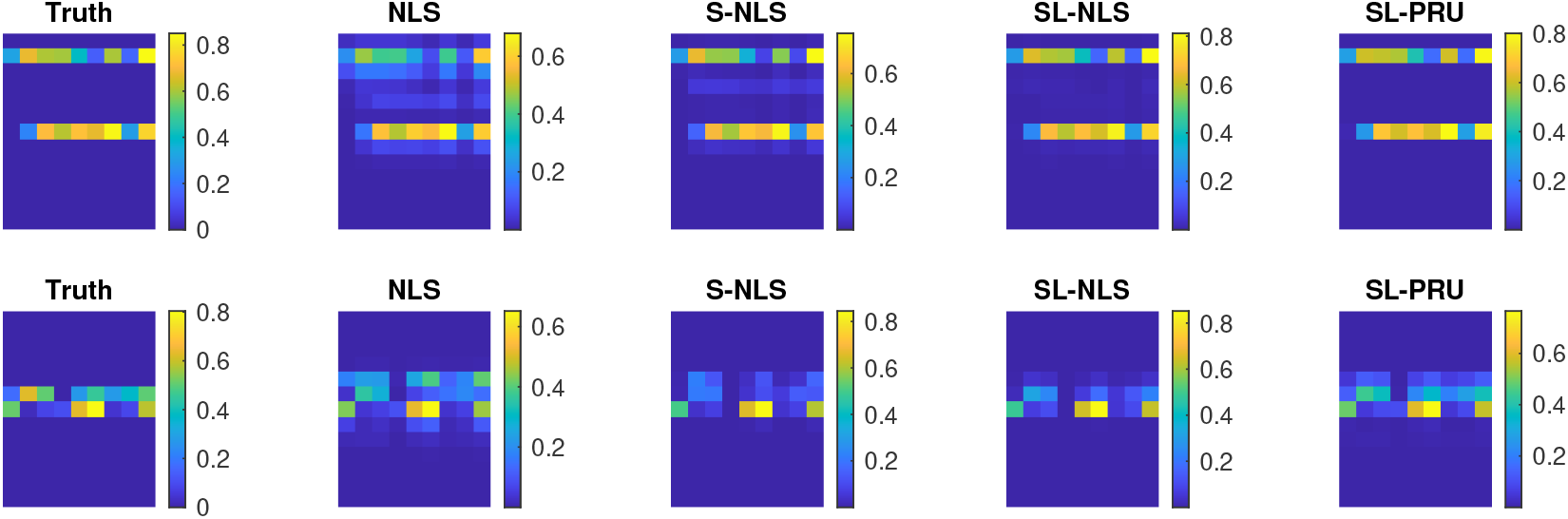
Graphical representation of simulated image pixels that contain two colocalized endmembers, either AF514 and RRX (highly uncorrelated endmembers, (Top row)) or AF555 and RRX (highly correlated endmembers (Bottom row)). In each row, the ‘‘Truth” matrix represents the ground truth starting simulation. Subsequent matrices represent the results of estimated abundances obtained from each of the unmixing methods that we considered. For each matrix, the 13 rows represent the 13 different endmembers used in the simulation and each column represents an independent pixel with varying intensity, scaled from 0-1. The color represents the mean abundance measure for each fluorophore from 1000 simulations of the same ground truth model after applying Poisson noise with SNR = 5 to each pixel.

From the reported experimental results in Figure 3 and the top row of Figure 5, it is observed that our proposed method can outperform other unmixing approaches under comparison here. For instance, as shown in Figure 3, our proposed method gives the lowest RMSEs with different SNRs. And as illustrated in Figure 5, the estimated average abundance matrix is more close to the true abundance matrix.

#### 3.2.2 Simulation 2: Correlated endmembers

We next apply SL-PRU to unmix simulated spectral images with correlated endmembers AF555 and RRX under the same experimental setup as in Simulation I. Similarly, the averaged RMSEs from 1, 000 repetitions of all the unmixing methods are plotted in Figure 4, and when SNR is set to 5, the average of the estimated abundance matrices for this scenario is plotted in the bottom row of Figure 5.

As shown in Figure 4 and the bottom row of Figure 5, SL-PRU also outperforms other unmixing approaches under comparison here. It is noted that the correlation between endmembers does have an impact on the performance of SL-PRU as well as other unmixing approaches, which coincides with our intuitive understanding of linear unmixing problems. However, the experiments carried out in our study also suggest that SL-PRU may outperform other unmixing approaches in unmixing spectral images with correlated endmembers, which could be an appealing feature of SL-PRU in some practical scenarios.

### 3.3 Unmixing real mixed biological images

We next apply SL-PRU to unmix a real biological image with known fluorophore labels, but unknown spatial distributions and abundances. A dental plaque smear hedgehog structure was obtained from a healthy volunteer via dental flossing and labeled in a FISH experiment with taxon-specific probes for 8 different genera or families of bacteria, with each probe conjugated to a different fluorescent reporter. Reference spectra were obtained from images of separate populations of E. coli cells labeled with FISH probes conjugated to the eight fluorophores used in the plaque smear experiment and imaged under identical acquisition settings. Reference spectra were extracted using our Poisson endmember extraction procedure. The values for sparseness and low-rank tuning parameters were selected using a heuristic approach. Visual inspection of the unmixed plaque smear images was performed over the range of tuning parameters described above and values that maximized expected cell morphologies and minimized salt and pepper noise in the image were chosen for the final unmixing result (Figure 6 and Supplementary Movie). To evaluate the performance of our SL-PRU on this real image, we performed a comparative, quantitative cellular morphological analysis against the unmixing result we obtained for the same image using the commercial microscope vendor’s linear unmixing algorithm. One plaque smear image set was used for quantitative comparison. The full z-stack image was unmixed in Zeiss Zen software, with endmember reference spectra extracted from single E. coli cells in images acquired with the same settings as the plaque smear image. The same spectral image data set was then unmixed using our SL-PRU approach. One central z-plane from both images was extracted and used for quantitative comparison. Unmixed images were imported into ImageJ. The Streptococcus and Veillonella channels were segmented using an intensity threshold determined algorithmically using the same algorithm, either “Otsu” or ‘Triangle” for the Streptococcus channels and Veillonella channels respectively. Circularity analysis was performed on each segmented image, defined as

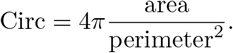

**Figure 6:**
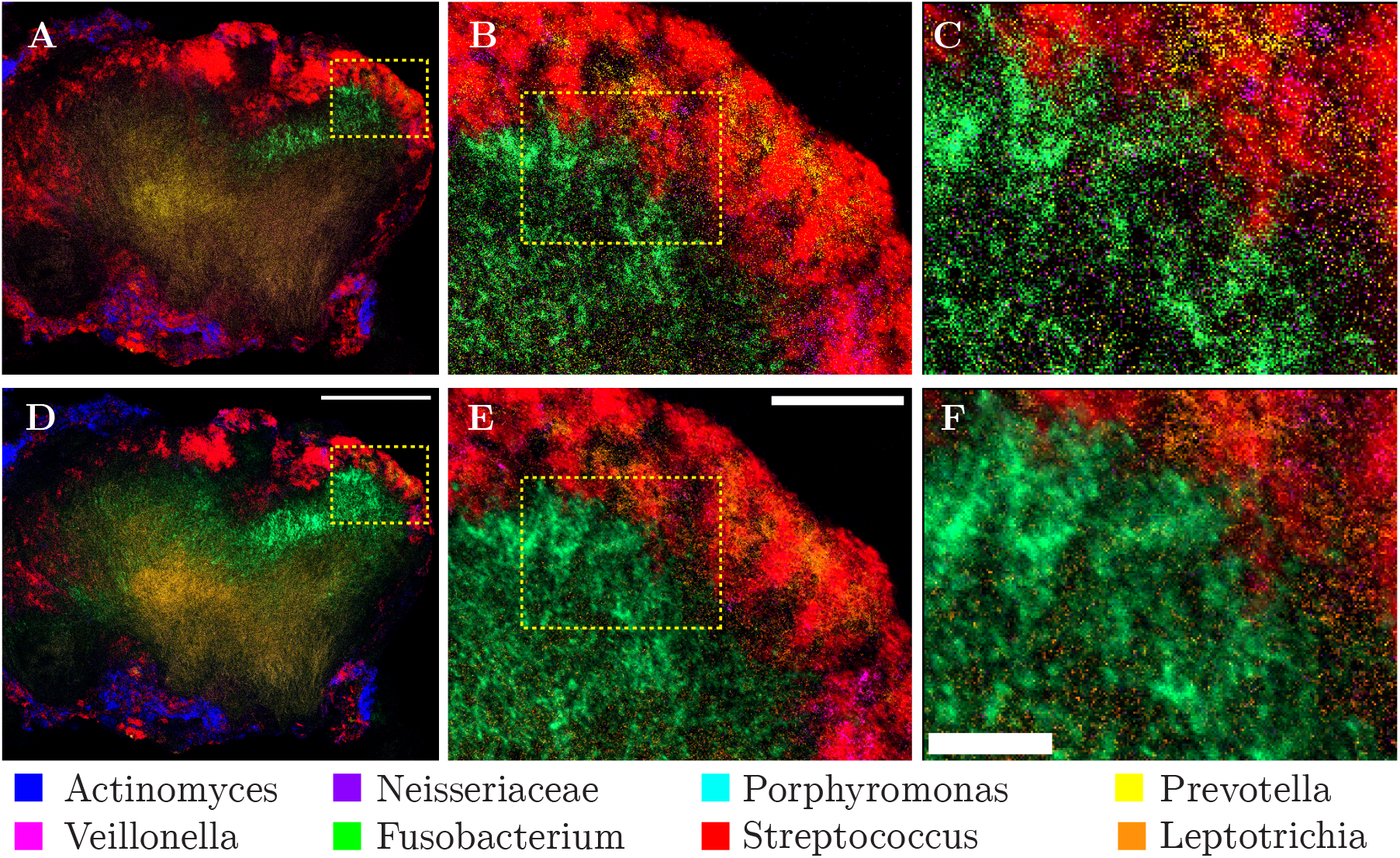
Qualitative comparison of least squares and SL-PRU unmixing on a real biological sample. A-F: Multi-spectral image of a dental plaque smear hedgehog structure after (A-C) least squares unmixing and (D-F) SL-PRU unmixing. Dashed boxes in (A) and (B) and in (C) and (D) indicate zoom area in C-D, and E-F respectively. Scale bars equal 100 μm (D), 25 μm, (E), 10 μm (F).

Streptococcus and Veillonella cells in the sample have characteristic, near-perfect spherical shapes. With this a priori information about these two labeled cells, we measured the circularity of these two cell populations in one central plane of the plaque smear image and found that SL-PRU improved the circularity measure by 16% for Streptococcus and 18.4% for Veillonella (Figure 7). Lastly, we performed a line scan analysis on the same region of interest in the Streptococcus channel in both unmixed images and found that the SL-PRU approach generated an image with less noise and more identifiable cell boundaries than the commercial vendor least squares unmixing (Supplementary Figure 2).

**Figure 7:**
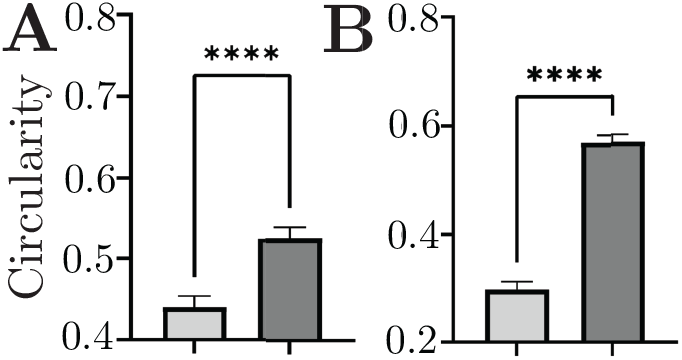
Quantitative comparison of mean circularity measurement per cell for two coccoid-shaped cells in the plaque structure: Streptococcus (A) and Veillonella (B). Light grey bars = results from least squares unmixing, dark grey bars = results from SL-PU. Error bars represent 95% confidence intervals. **** = *p* ! 0.001 Welch’s *t*-test.

## 4 Discussion

Multispectral imaging has allowed the visualization and quantification of large numbers of targets simultaneously within biological specimens. The linear model assumption behind spectral unmixing holds for most biological images acquired from specimens that are thin and therefore have minimal scattering effects and in which fluorescent tags label components, e.g., cells or macromolecules that are well separated in space, beyond the minimal Förster Resonance Energy Transfer distance. The least squares approach to unmixing is essentially a multi-output regression problem, which assumes a Gaussian noise distribution in the recorded signals; while it is known that the dominant noise source in fluorescence images follows a Poisson distribution. To better accommodate such a Poisson distribution, and to improve fidelity in fluorophore abundance estimation in the unmixing process by using prior information about our labeled samples, we have developed in this paper a sparse and low-rank Poisson (SL-PRU) approach to multispectral unmixing. Specifically, we considered a two-step approach where we first extract endmember information through reference images using unregularized Poisson regression and then learn the abundance information via the proposed regularized Poisson approach. In the case of multiplex labeled cells such as the microbial biofilm samples used here, we assume that while many dozens of fluorophores might be present in the sample, for any individual pixel, the number of fluorophores present approaches one. We demonstrated the effectiveness of the proposed approach through experimental results on model images and real biological samples reported above.

We believe that our SL-PRU unmixing approach will be generally applicable to a wide variety of image datasets of multi-plex, fluorescently labeled cells. As microscope detector technologies improve, and the need for extracting information from ever lower numbers of photons increases, the dominant noise source in fluorescence images of cells is expected to shift ever more toward the Poisson-distributed, physically unavoidable photon shot noise. Even as the number of labeled targets increases, the physical constraints of these targets, whether they be cells or macromolecules, limits their simultaneous occurrence in pixels in digitally recorded images, although we recognize that the finite resolution of the light microscope and the labeling of target molecules that are sufficiently small and diffusible together put limitations on the sparsity assumption in some situations. In implementing our low-rank penalty term, we used a sliding window approach and restricted our neighborhood size to 3 x 3 pixels. In general, the size of the sliding window should be dictated by the prior information available about the sample, e.g., the relative size of the labeled targets vs pixel dimensions in the image.

Finally, the effectiveness of the proposed sparse and low-rank Poisson approach has justified the importance of the low-rankness constraint in abundance estimation, which indicates the similarity of abundances of neighboring pixels. Note that such similarities, which agree with our intuitive understanding of spectral images, can be used to account for the spatial information among neighboring pixels. Recall that spectral images are three-way tensors and we unfold these tensor data into matrices before unmixing them. Therefore, further improved results may be obtained if the spatial information of the original tensor spectral data is taken into account in unmixing.

## 5 Conclusion

The existence of photon shot noise in biological fluorescence spectral images motivates us to address the unmixing problem through a Poisson approach instead of NLS. We also incorporate the spatial information by imposing sparsity and low-rankness constraints in a localized pattern. The unmixing results from both simulated data and real-world biological images demonstrate that our proposed approach SL-PRU can identify endmembers and estimate the corresponding abundances with increased accuracy over existing unmixing approaches.

## Acknowledgement

This work was supported by the National Science Foundation [DMS-2111080 to Y.F., DMS-2110836, IIS-2103450, and IIS-2110546 to Y.Y.]; the Simons Foundation [#572064 to Y.F.]; and the National Institutes of Health [R01DE030927 to A.M.V.].

