## Supplementary Methods and Data for "Unmixing Biological Fluorescence Image Data with Sparse and Low-Rank Poisson Regression"

#### Contents

**1 The Proposed Algorithms for Solving PNMF and SL-PRU** i

**2 Supplementary Table 1: Salient Characteristics of FISH Probes** vi

**3 Supplementary Figure 1: Emission Spectra of Endmembers and the Extracted Endmembers** vii

**4 Supplementary Table 2: Descriptions of Fluorophores** viii

**5 Supplementary Figure 2: Quantitative Line Scan Analysis on Dental Plaque Smear Images** ix

**6 Supplementary Movie: SL-RPU Unmixed Dental Plaque Smear** x

**References** xi

#### 1 The Proposed Algorithms for Solving PNMF and SL-PRU

We first present the developed algorithms for solving the endmember extraction problem through PNMF and for solving the abundance estimation problem through SL-PRU, respectively. To this end, we first recall that the endmember $\mathbf{m} \in \mathbb{R}_+^C$ extracted from a reference image $\mathbf{Y_m} \in \mathbb{R}_+^{C\times N}$ and the corresponding abundances $\mathbf{a} \in \mathbb{R}_+^N$ are obtained through Poisson Nonnegative Matrix Factorization (PNMF):

$$\min_{\mathbf{m}\in\mathbb{R}_+^C,\mathbf{a}\in\mathbb{R}_+^N} \mathbf{1}_C^\top \left[\mathbf{m}\mathbf{a}^\top - \mathbf{Y_m} \circ \log\left(\mathbf{m}\mathbf{a}^\top\right)\right] \mathbf{1}_N, \tag{1}$$

where $\mathbf{1}_C$ and $\mathbf{1}_N$ are vectors of length $C$ and $N$ whose entries are all 1 and $\circ$ denotes element-wise multiplication. PNMF (1) can be solved by *multiplicative update algorithm* (Lee and Seung, 2000) which is a diagonally rescaled version of gradient descent. Denoting any variable $\mathbf{X}$ at the $t$-th, iteration as $\mathbf{X}^{(t)}$ and the maximum norm as $\|\cdot\|_\infty$, the pseudocode of the multiplicative update algorithm is provided in Algorithm 1. The abundance vector $\mathbf{a}$ and the endmember vector $\mathbf{m}$ are initialized with random values that follow the standard uniform distribution $U(0,1)$. At each iteration, the endmember vector is standardized by dividing by its maximum for uniqueness. The algorithm stops when the relative change of the standardized endmember $\widetilde{\mathbf{m}}$ between the $(t-1)$-th and the $t$-th iterations given by

$$\frac{\|\widetilde{\mathbf{m}}^{(t)} - \widetilde{\mathbf{m}}^{(t-1)}\|_2^2}{\|\widetilde{\mathbf{m}}^{(t-1)}\|_2^2}$$

is less than a small threshold value.

**Input:** A reference image $\mathbf{Y_m} \in \mathbb{R}_+^{C\times N}$;
**Output:** The standardized endmember vector $\widetilde{\mathbf{m}}$;
**Initialization:** $\mathbf{a}^{(0)}$, $\mathbf{m}^{(0)}$ ;
**repeat**

$a_n^{(t)} \leftarrow \widetilde{a}_n^{(t-1)} \cdot \left(\frac{\sum_c y_{cn}\widetilde{m}_c^{(t-1)}}{\widetilde{m}_c^{(t-1)}\widetilde{a}_n^{(t-1)}}\right) \Big/ \left(\sum_c \widetilde{m}_c^{(t-1)}\right)$;

$m_c^{(t)} \leftarrow \widetilde{m}_c^{(t-1)} \cdot \left(\frac{\sum_n y_{cn} a_n^{(t)}}{\widetilde{m}_c^{(t-1)} a_n^{(t)}}\right) \Big/ \left(\sum_n a_n^{(t)}\right)$;

$\widetilde{m}_c^{(t)} \leftarrow m_c^{(t)} / \|\mathbf{m}^{(t)}\|_\infty$;

$\widetilde{a}_n^{(t)} \leftarrow a_n^{(t)} \cdot \|\mathbf{m}^{(t)}\|_\infty$;

**until** *the stopping criterion is satisfied*;

**Algorithm 1:** Multiplicative update algorithm for PNMF (1)

We are now in a position to detail the proposed algorithm for solving the proposed regularized sparse and low-rank Poisson regression unmixing approach (SL-PRU) that reads

$$\min_{\mathbf{A}\in\mathbb{R}_+^{R\times N}} \mathbf{1}_C^\top \left[\mathbf{MA} - \mathbf{Y} \circ \log\left(\mathbf{MA}\right)\right] \mathbf{1}_N + \lambda_1 \|\mathbf{A}\|_{\mathbf{w}_p,*} + \lambda_2 \|\mathbf{W}_q \mathbf{A}\|_{2,1}. \tag{2}$$

Inspired by the work in Giampouras *et al.* (2016), an alternating direction method of multipliers (ADMM) technique (Boyd *et al.*, 2011) is adopted in our study by first letting all elements of $\mathbf{w}_p$ be equal to ensure the convexity of the low-rankness regularization term in SL-PRU (2). Similar to the work in Giampouras *et al.* (2016), we introduce auxiliary variables $\mathbf{V}_1 \in \mathbb{R}^{C\times N}$, $\mathbf{V}_2, \mathbf{V}_3, \mathbf{V}_4 \in \mathbb{R}^{R\times N}$ and reformulate SL-PRU (2) as follows

$$\begin{aligned} \min_{\mathbf{U},\mathbf{V}_1,\mathbf{V}_2,\mathbf{V}_3,\mathbf{V}_4} \quad & \mathbf{1}_C^\top \left[\mathbf{V}_1 - \mathbf{Y} \circ \log\left(\mathbf{V}_1\right)\right] \mathbf{1}_N + \lambda_1 \|\mathbf{V}_2\|_{\mathbf{w}_p,*} + \lambda_2 \|\mathbf{W}_q \mathbf{V}_3\|_{2,1} + \mathcal{I}_{\mathbb{R}_+}(\mathbf{V}_4), \\ \text{s.t.} \quad & \mathbf{V}_1 = \mathbf{MU}, \mathbf{V}_2 = \mathbf{U}, \mathbf{V}_3 = \mathbf{U}, \mathbf{V}_4 = \mathbf{U}, \end{aligned} \tag{3}$$

where $\mathcal{I}_{\mathbb{R}_+}(\cdot)$ is the indicator function which is zero if all the entries are nonnegative and infinity otherwise. The augmented Lagrangian function for the constrained optimization problem (3) is given as follows:

$$\begin{aligned} &\mathcal{L}_1(\mathbf{U}, \mathbf{V}_1, \mathbf{V}_2, \mathbf{V}_3, \mathbf{V}_4, \mathbf{D}_1, \mathbf{D}_2, \mathbf{D}_3, \mathbf{D}_4) \\ &= \mathbf{1}_C^\top \left[\mathbf{V}_1 - \mathbf{Y} \circ \log\left(\mathbf{V}_1\right)\right] \mathbf{1}_N + \lambda_1 \|\mathbf{V}_2\|_{\mathbf{w}_p,*} + \lambda_2 \|\mathbf{W}_q \mathbf{V}_3\|_{2,1} + \mathcal{I}_{\mathbb{R}_+}(\mathbf{V}_4) \\ &\quad + \mathrm{tr}(\mathbf{D}_1^\top(\mathbf{V}_1 - \mathbf{MU})) + \mathrm{tr}(\mathbf{D}_2^\top(\mathbf{V}_2 - \mathbf{U})) + \mathrm{tr}(\mathbf{D}_3^\top(\mathbf{V}_3 - \mathbf{U})) + \mathrm{tr}(\mathbf{D}_4^\top(\mathbf{V}_4 - \mathbf{U})) \\ &\quad + \frac{\mu}{2}\left(\|\mathbf{MU} - \mathbf{V}_1\|_F^2 + \|\mathbf{U} - \mathbf{V}_2\|_F^2 + \|\mathbf{U} - \mathbf{V}_3\|_F^2 + \|\mathbf{U} - \mathbf{V}_4\|_F^2\right), \end{aligned} \tag{4}$$

where $\mathbf{D}_1 \in \mathbb{R}^{C\times N}$, $\mathbf{D}_2, \mathbf{D}_3, \mathbf{D}_4 \in \mathbb{R}^{R\times N}$ denote the Lagrange multipliers, $\mathrm{tr}(\cdot)$ denotes matrix trace, $\mu > 0$ is a Lagrange multiplier regularization parameter, and $\|\cdot\|_F$ denotes the Frobenius norm.

Denoting the identity matrix of size $k \times k$ as $\mathbf{I}_k$ and the scaled Lagrange multipliers as $\mathbf{D}'_i = \mathbf{D}_i/\mu, i = 1, 2, 3, 4$, the augmented Lagrangian function $\mathcal{L}_1$ can be rewritten as

$$\begin{aligned}\mathcal{L}_2(\mathbf{U}, \mathbf{V}, \mathbf{D}) = &\mathbf{1}_C^\top \left[\mathbf{V}_1 - \mathbf{Y} \circ \log\left(\mathbf{V}_1\right)\right] \mathbf{1}_N + \lambda_1 \|\mathbf{V}_2\|_{\mathbf{w}_p,*} + \lambda_2 \|\mathbf{W}_q \mathbf{V}_3\|_{2,1} + \mathcal{I}_{\mathbb{R}_+}(\mathbf{V}_4) \\ &+ \frac{\mu}{2} \|\mathbf{G}\mathbf{U} + \mathbf{B}\mathbf{V} - \mathbf{D}\|_F^2,\end{aligned} \tag{5}$$

where

$$\mathbf{V} = \begin{bmatrix}\mathbf{V}_1 \\ \mathbf{V}_2 \\ \mathbf{V}_3 \\ \mathbf{V}_4\end{bmatrix}, \mathbf{D} = \begin{bmatrix}\mathbf{D}'_1 \\ \mathbf{D}'_2 \\ \mathbf{D}'_3 \\ \mathbf{D}'_4\end{bmatrix}, \mathbf{G} = \begin{bmatrix}\mathbf{M} \\ \mathbf{I}_R \\ \mathbf{I}_R \\ \mathbf{I}_R\end{bmatrix}, \mathbf{B} = -\mathbf{I}_{C+3R}.$$

The proposed ADMM-type algorithm for solving SL-PRU sequentially optimizes (4) or (5) with respect to each variable while the other variables remain as the latest values. Note that the augmented Lagrangian (5) is convex w.r.t. $\mathbf{U}$, $\mathbf{V}_1$, $\mathbf{V}_2$, $\mathbf{V}_3$, and $\mathbf{V}_4$, respectively, due to the assumption that all elements of $\mathbf{w}_p$ are equal and the fact that all entries of $\mathbf{W}_q$ are nonnegative. Therefore, at the $t$-th iteration, the updates of the abundance matrix $\mathbf{U}$ and the auxiliary variables $\mathbf{V}_1$, $\mathbf{V}_2$, $\mathbf{V}_3$, and $\mathbf{V}_4$ can be deduced, respectively, as follows:

**Updating $\mathbf{U}$**: The minimization of $\mathcal{L}_2$ w.r.t. $\mathbf{U}$ at the $t$-th iteration is equivalent to

$$\frac{\partial \frac{\mu}{2} \|\mathbf{G}\mathbf{U} + \mathbf{B}\mathbf{V}^{(t-1)} - \mathbf{D}\|_F^2}{\partial \mathbf{U}} = 0,$$

which yields

$$\mu \mathbf{G}^\top (\mathbf{G}\mathbf{U} + \mathbf{B}\mathbf{V} - \mathbf{D}) = \mu \left[\left(\mathbf{M}^\top \mathbf{M} + 3\mathbf{I}_R\right) \mathbf{U} - \mathbf{G}^\top (\mathbf{B}\mathbf{V} - \mathbf{D})\right] = 0.$$

As a result, we have

$$\begin{aligned}\mathbf{U}^{(t)} = &\arg\min_{\mathbf{U}} \mathcal{L}_2\left(\mathbf{U}, \mathbf{V}^{(t-1)}, \mathbf{D}^{(t-1)}\right) = \left(\mathbf{M}^\top \mathbf{M} + 3\mathbf{I}_R\right)^{-1} \\ &\left[\mathbf{M}^\top \left(\mathbf{V}_1^{(t-1)} + \mathbf{D}_1^{'(t-1)}\right) + \mathbf{V}_2^{(t-1)} + \mathbf{D}_2^{'(t-1)} + \mathbf{V}_3^{(t-1)} + \mathbf{D}_3^{'(t-1)} + \mathbf{V}_4^{(t-1)} + \mathbf{D}_4^{'(t-1)}\right].\end{aligned}$$

**Updating $\mathbf{V}_1$**: Denoting the $(c, n)$-th entry of $\mathbf{V}_1$ as $v_{cn}$ with $c = 1, \cdots, C$ and $n = 1, \cdots, N$, we have

$$\mathbf{1}_C^\top \left[\mathbf{V}_1 - \mathbf{Y} \circ \log\left(\mathbf{V}_1\right)\right] \mathbf{1}_N = \sum_{c,n} [v_{cn} - y_{cn} \log(v_{cn})].$$

Since

$$\left(\frac{\partial \mathbf{1}_C^\top \left[\mathbf{V}_1 - \mathbf{Y} \circ \log\left(\mathbf{V}_1\right)\right] \mathbf{1}_N}{\partial \mathbf{V}_1}\right)_{cn} = \frac{\partial [v_{cn} - y_{cn} \log(v_{cn})]}{\partial v_{cn}} = 1 - \frac{y_{cn}}{v_{cn}},$$

and

$$\left(\frac{\partial \frac{\mu}{2} \|\mathbf{M}\mathbf{U}^{(t)} - \mathbf{V}_1 - \mathbf{D}'_1\|_F^2}{\partial \mathbf{V}_1}\right)_{cn} = \mu [v_{cn} + (\mathbf{D}_1^{'(t-1)} - \mathbf{M}\mathbf{U}^{(t)})_{cn}],$$

we know that minimizing $\mathcal{L}_2$ w.r.t. $v_{cn}$ is equivalent to

$$v_{cn}^2 - (\mathbf{V}_1'^{(t)})_{cn} \cdot v_{cn} - \frac{y_{cn}}{\mu} = 0,$$

where $\mathbf{V}_1'^{(t)} = \mathbf{M}\mathbf{U}^{(t)} - \mathbf{D}_1^{'(t-1)} - 1/\mu$. Solving the above quadratic equation w.r.t. $v_{cn}$, we have

$$\mathbf{V}_1^{(t)} = \arg\min_{\mathbf{V}_1} \mathcal{L}_2 \left( \mathbf{U}^{(t)}, \begin{bmatrix} \mathbf{V}_1 \\ \mathbf{V}_2^{(t-1)} \\ \mathbf{V}_3^{(t-1)} \\ \mathbf{V}_4^{(t-1)} \end{bmatrix}, \mathbf{D}^{(t-1)} \right) = \frac{\mathbf{V}_1'^{(t)} + \sqrt{\mathbf{V}_1'^{(t)} \circ \mathbf{V}_1'^{(t)} + 4\mathbf{Y}/\mu}}{2},$$

where $\sqrt{\cdot}$ denotes the element-wise square root.

**Updating $\mathbf{V}_2$**: Minimizing $\mathcal{L}_2$ w.r.t. $\mathbf{V}_2$ is equivalent to

$$\min_{\mathbf{V}_2} \lambda_1 \|\mathbf{V}_2\|_{\mathbf{w}_p,*} + \frac{\mu}{2} \|\mathbf{U} - \mathbf{V}_2 - \mathbf{D}_2'\|_F^2,$$

which can be solved by a soft-thresholding operation (Cai *et al.*, 2010) on the singular values of $\mathbf{V}_2$. Recall that the singular value decomposition of $\mathbf{U}^{(t)} - \mathbf{D}_2^{'(t-1)}$ is $\mathbf{S}_l \mathbf{\Sigma}^{(t)} \mathbf{S}_r^\top$, the soft-thresholding function on each diagonal element of $\mathbf{\Sigma}^{(t)}$, i.e., $\sigma_p^{(t)}$, with parameter $\lambda_1 \mathbf{w}_p/\mu$ is

$$\max\{\mathbf{0}, \sigma_p^{(t)} - \lambda_1 \mathbf{w}_p/\mu\},$$

for any $p = 1, 2, \cdots, \mathrm{rank}(\mathbf{V}_2)$. Thus, the optimization w.r.t. $\mathbf{V}_2$ gives

$$\mathbf{V}_2^{(t)} = \arg\min_{\mathbf{V}_2} \mathcal{L}_2 \left( \mathbf{U}^{(t)}, \begin{bmatrix} \mathbf{V}_1^{(t)} \\ \mathbf{V}_2 \\ \mathbf{V}_3^{(t-1)} \\ \mathbf{V}_4^{(t-1)} \end{bmatrix}, \mathbf{D}^{(t-1)} \right) = \mathbf{S}_l[\mathrm{sign}(\mathbf{\Sigma}^{(t)}) \circ \max\{\mathbf{0}, \mathbf{\Sigma}^{(t)} - \lambda_1 \mathrm{diag}(\mathbf{w}_p)/\mu\}]\mathbf{S}_r^\top,$$

where $\mathrm{sign}(\cdot)$ is the element-wise sign function, $\max\{\cdot,\cdot\}$ denotes the element-wise max function, and $\mathrm{diag}(\cdot)$ creates a matrix with diagonal elements equal to the vector elements.

**Updating $\mathbf{V}_3$**: Minimizing $\mathcal{L}_2$ w.r.t. $\mathbf{V}_3$ is equivalent to

$$\min_{\mathbf{V}_3} \lambda_2 \|\mathbf{W}_q \mathbf{V}_3\|_{2,1} + \frac{\mu}{2} \|\mathbf{U} - \mathbf{V}_3 - \mathbf{D}_3'\|_F^2,$$

which can be solved by a vectorial soft-thresholding operation (Wright *et al.*, 2009) on each row of $\mathbf{V}_3^{(t)}$. More specifically, denoting $\mathbf{V}_{3,r}^{(t)}$ as the $r$-th row of $\mathbf{V}_3^{(t)}$ and $\mathbf{x}_r^{(t)}$ as the $r$-th row of $\mathbf{U}^{(t)} - \mathbf{D}_3^{'(t-1)}$ where $r = 1, \ldots, R$, each row of $\mathbf{V}_3$ is updated sequentially as

$$\mathbf{V}_{3,r}^{(t)} = \arg\min_{\mathbf{V}_{3,r}} \mathcal{L}_2 \left( \mathbf{U}^{(t)}, \begin{bmatrix} \mathbf{V}_1^{(t)} \\ \mathbf{V}_2^{(t)} \\ \mathbf{V}_{3,1}^{(t)} \\ \vdots \\ \mathbf{V}_{3,r-1}^{(t)} \\ \mathbf{V}_{3,r} \\ \mathbf{V}_{3,r+1}^{(t-1)} \\ \vdots \\ \mathbf{V}_{3,R}^{(t-1)} \\ \mathbf{V}_4^{(t-1)} \end{bmatrix}, \mathbf{D}^{(t-1)} \right) = \frac{\mathbf{x}_r^{(t)} \max\{\|\mathbf{x}_r^{(t)}\|_2 - \lambda_2 w_{q,r}/\mu, 0\}}{\max\{\|\mathbf{x}_r^{(t)}\|_2 - \lambda_2 w_{q,r}/\mu, 0\} + \lambda_2 w_{q,r}/\mu}.$$

**Input:** The data matrix $\mathbf{Y} \in \mathbb{R}_{+}^{C\times N}$, and the endmember matrix $\mathbf{M} \in \mathbb{R}_{+}^{C\times R}$;
**Output:** The abundance matrix $\mathbf{U}$;
**Initialization:** $\mathbf{U}^0, \mathbf{V}_i^0, \mathbf{D}_i^0, \quad i = 1, 2, 3, 4$ ;
**repeat**

$\mathbf{U}^{(t)} \leftarrow \left(\mathbf{M}^\top\mathbf{M} + 3\mathbf{I}_R\right)^{-1} \Big[\mathbf{M}^\top \left(\mathbf{V}_1^{(t-1)} + \mathbf{D}_1^{'(t-1)}\right) + \mathbf{V}_2^{(t-1)} + \mathbf{D}_2^{'(t-1)} + \mathbf{V}_3^{(t-1)} + \mathbf{D}_3^{'(t-1)} + \mathbf{V}_4^{(t-1)} + \mathbf{D}_4^{'(t-1)}\Big]$;

$\mathbf{V}_1^{(t)} \leftarrow \left(\mathbf{V}_1^{(t)} + \sqrt{\mathbf{V}_1^{(t)} \circ \mathbf{V}_1^{(t)} + 4\mathbf{Y}/\mu}\right) \Big/ 2$;

$\mathbf{V}_2^{(t)} \leftarrow \mathbf{S}_l[\mathrm{sign}(\boldsymbol{\Sigma}) \circ \max\{\mathbf{0}, \boldsymbol{\Sigma} - \lambda_1 \mathrm{diag}(\mathbf{w}_p)/\mu\}]\mathbf{S}_r^\top$;

$\mathbf{V}_{3,r}^{(t)} \leftarrow (\mathbf{x}_r^{(t)} \max\{\|\mathbf{x}_r^{(t)}\|_2 - \lambda_2 w_{q,r}/\mu, 0\})/(\max\{\|\mathbf{x}_r^{(t)}\|_2 - \lambda_2 w_{q,r}/\mu, 0\} + \lambda_2 w_{q,r}/\mu)$, $r = 1, \ldots, R$;

$\mathbf{V}_4^{(t)} \leftarrow \max\left\{\mathbf{U}^{(t)} - \mathbf{D}_4^{\prime(t-1)}, \mathbf{0}\right\}$;

$\mathbf{D}_1^{\prime(t)} \leftarrow \mathbf{D}_1^{\prime(t-1)} - \mathbf{M}\mathbf{U}^{(t)} + \mathbf{V}_1^{(t)}$;

$\mathbf{D}_i^{\prime(t)} \leftarrow \mathbf{D}_i^{\prime(t-1)} - \mathbf{U}^{(t)} + \mathbf{V}_i^{(t)}, \quad i = 2, 3, 4$;

**until** *the stopping criteria are satisfied*;

**Algorithm 2:** The proposed ADMM-type algorithm for SL-PRU (2)

**Updating $\mathbf{V}_4$**: Minimizing $\mathcal{L}_2$ w.r.t. $\mathbf{V}_4$ is equivalent to

$$\min_{\mathbf{V}_4} \mathcal{I}_{\mathbb{R}_+}(\mathbf{V}_4) + \frac{\mu}{2}\|\mathbf{U} - \mathbf{V}_4 - \mathbf{D}_4'\|_F^2,$$

where the first term is to project $\mathbf{V}_4$ onto the nonnegative orthant. Thus, the optimization w.r.t. $\mathbf{V}_4$ gives

$$\mathbf{V}_4^{(t)} = \arg\min_{\mathbf{V}_4} \mathcal{L}_2 \left( \mathbf{U}^{(t)}, \begin{bmatrix} \mathbf{V}_1^{(t)} \\ \mathbf{V}_2^{(t)} \\ \mathbf{V}_3^{(t)} \\ \mathbf{V}_4 \end{bmatrix}, \mathbf{D}^{(t-1)} \right) = \max\left\{\mathbf{U}^{(t)} - \mathbf{D}_4^{'(t-1)}, \mathbf{0}\right\}.$$

**Updating $\mathbf{D}_1'$, $\mathbf{D}_2'$, $\mathbf{D}_3'$, and $\mathbf{D}_4'$**: These scaled Lagrange multipliers are updated as follows

$$\mathbf{D}_1^{\prime(t)} = \mathbf{D}_1^{\prime(t-1)} - \mathbf{M}\mathbf{U}^{(t)} + \mathbf{V}_1^{(t)}, \quad \mathbf{D}_i^{\prime(t)} = \mathbf{D}_i^{\prime(t-1)} - \mathbf{U}^{(t)} + \mathbf{V}_i^{(t)}, \quad i = 2, 3, 4.$$

The stopping criteria adopted in the algorithm are based on the primal and dual residuals (Boyd *et al.*, 2011) $\mathbf{r}_p$ and $\mathbf{r}_d$ given by

$$\mathbf{r}_p = \mathbf{G}\mathbf{U}^{(t)} + \mathbf{B}\mathbf{V}^{(t)}, \quad \mathbf{r}_d = \mu\mathbf{G}^\top\mathbf{B}\left(\mathbf{V}^{(t)} - \mathbf{V}^{(t-1)}\right),$$

that go to 0, respectively, as $t \to \infty$. The algorithm terminates whenever any of the $\ell_2$ norms of $\mathbf{r}_p$ or $\mathbf{r}_d$ is less than a small threshold value or some number of iterations is reached. To enhance the performance of the algorithm, as done in Giampouras *et al.* (2016), we also update the weights $\mathbf{w}_p$ and $\mathbf{W}_q$ based on $\mathbf{U}^{(t)}$ at the $t$-th iteration as follows

$$w_{p,i}^{(t)} = \frac{1}{\sigma_i(\mathbf{U}^{(t)}) + \varepsilon}, \quad w_{q,r} = \frac{1}{\|\mathbf{u}_r^{(t)}\|_2 + \varepsilon},$$

where $\mathbf{u}_r^{(t)}$ denotes the $r$-th row of $\mathbf{U}^{(t)}$ and $\varepsilon > 0$ is assigned a small value to avoid singularities.

The pseudo-code of the proposed algorithm for solving SL-PRU is presented in Algorithm 2.

### 2 Supplementary Table 1: Salient Characteristics of FISH Probes

| Probe name | Target | Sequence | Fluorophore in plaque smear image | Reference |
|---|---|---|---|---|
| ACT-476 | Actinomyces | ATCCAGCTACCGTCAACC | Alexafluor 488 | (Gmür and Lüthi-Schaller, 2007) |
| STR-405 | Streptococcus | TAGCCGTCCCTTTCTGGT | Alexafluor 594 | (Paster, Bruce J *et al.*, 1998) |
| FUS-714 | Fusobacterium | GGCTTCCCCATCGGCATT | Alexafluor 555 | (Valm, Alex M *et al.*, 2011) |
| LEP-568 | Leptotrichia | GCCTAGATGCCCTTTATG | Alexafluor 660 | (Valm, Alex M *et al.*, 2011) |
| NEI-1030 | Neisseriaceae | CCTGTGTTACGGCTCCCG | Tetrachlorofluorescein | (Valm, Alex M *et al.*, 2011) |
| PGI-350 | Porphyromonas | CCTCACGCCTTACGACGG | Alexafluor 647 | (Valm, Alex M *et al.*, 2011) |
| VEI-488 | Veillonella | CCGTGGCTTTCTATTCCG | Alexafluor 514 | (Chalmers *et al.*, 2008) |
| PRV-392 | Prevotella | GCACGCTACTTGGCTGG | Alexafluor 633 | (Diaz *et al.*, 2006) |
| PAS-111 | Pasteurellaceae | TCCCAAGCATTACTCACC | Rhodamine Red-X | (Valm, Alex M *et al.*, 2011) |
| EUB-338 | All bacteria | GCTGCCTCCCGTAGGAGT | Used for E. coli reference standards | (Amann *et al.*, 1990) |

Supplementary Table 1: Salient characteristics of all FISH probes used in this study

### 3 Supplementary Figure 1: Emission Spectra of Endmembers and the Extracted Endmembers

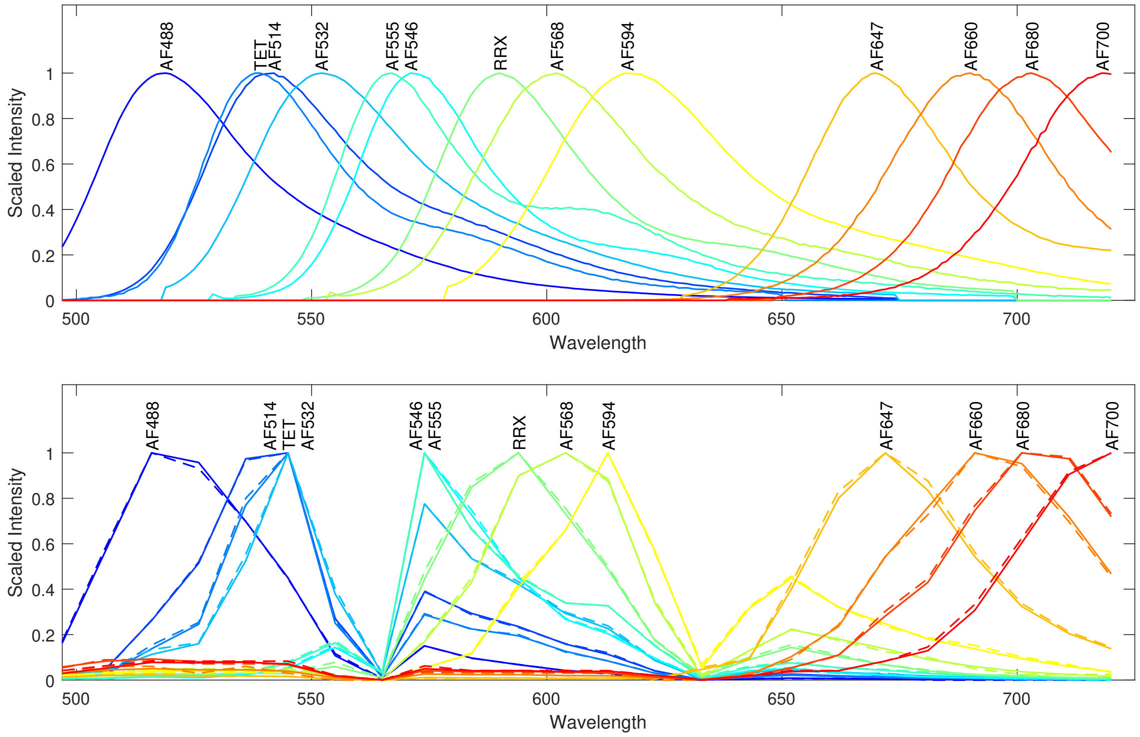


Supplementary Figure 1: Empirically measured spectra of thirteen endmembers from Supplementary Table 2 above. Top row: Fluorometer data provided by the manufacturers of the dyes sampled in 1 nm wavelength bands. Bottom row: Endmembers extracted by arithmetic mean (dashed curves) method and PNMF (solid curves) from real images of labeled E. coli acquired on a spectral confocal microscope with 9.8 nm wavelength bands.

### 4 Supplementary Table 2: Descriptions of Fluorophores

| Fluorophore | Abbreviation | Peak Excitation $\lambda$ (nm) | Peak Emission $\lambda$ (nm) |
|---|---|---|---|
| Alexafluor 488 | AF488 | 495 | 519 |
| Alexafluor 514 | AF514 | 517 | 542 |
| Tetrachlorofluorescein | TET | 522 | 539 |
| Alexafluor 532 | AF532 | 532 | 553 |
| Alexafluor 546 | AF546 | 556 | 573 |
| Alexafluor 555 | AF555 | 555 | 565 |
| Rhodamine Red-X | RRX | 560 | 580 |
| Alexafluor 568 | AF568 | 578 | 603 |
| Alexafluor 633 | AF633 | 621 | 639 |
| Alexafluor 647 | AF647 | 650 | 665 |
| Alexafluor 660 | AF660 | 663 | 690 |
| Alexafluor 680 | AF680 | 679 | 702 |
| Alexafluor 700 | AF700 | 702 | 723 |

Supplementary Table 2: Names, abbreviations, and peak excitation and emission wavelengths for all fluorophores used in this study.

### 5 Supplementary Figure 2: Quantitative Line Scan Analysis on Dental Plaque Smear Images

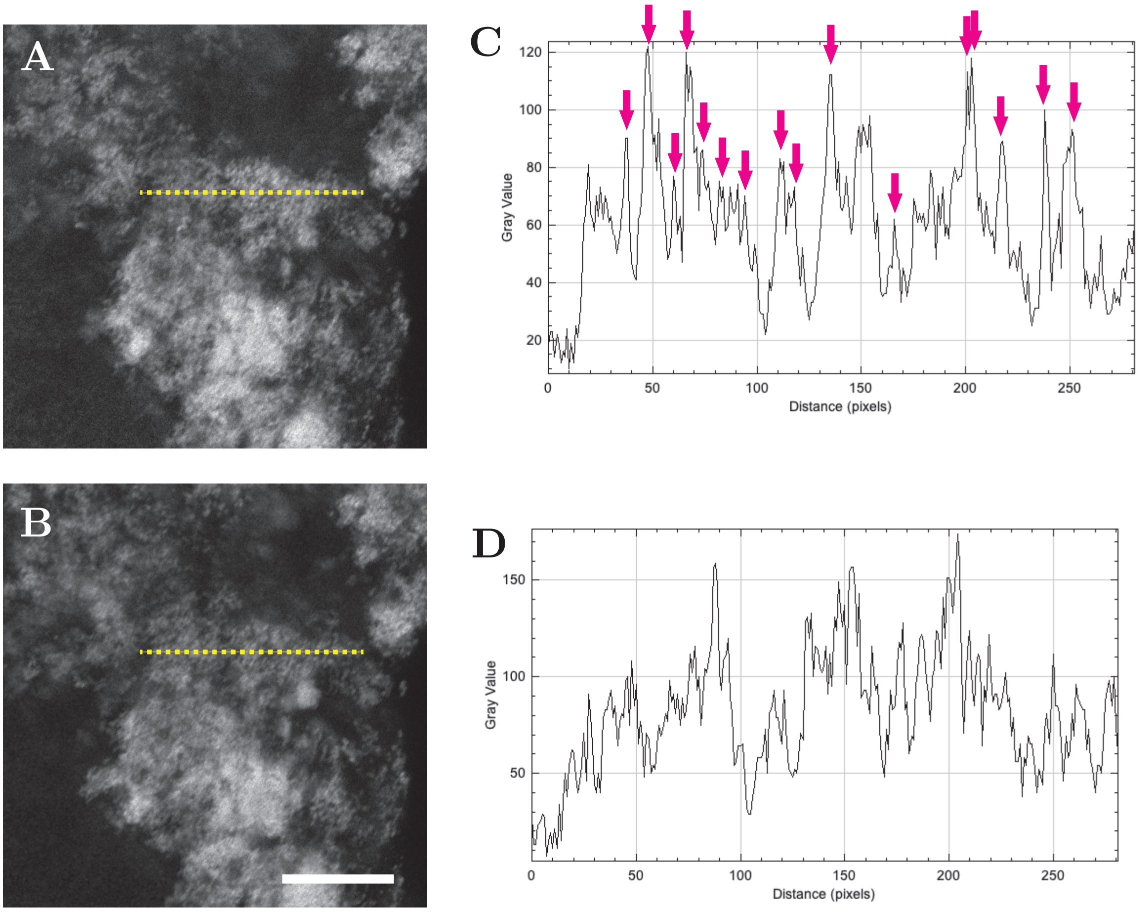


Supplementary Figure 2: A-B. Region of interest images from the Streptococcus (Alexa-fluor 594) channel from (A) SL-PRU unmixed image and (B) commercial least squares unmixed image. Yellow dotted lines show where line scan analysis was performed in each image. Bar = 25 µm. C-D: Line scan analysis results (intensity vs. pixel position) for SL-PRU unmixing (C) and least squares unmixing (D) along the length of the lines in A & B. Magenta arrows in (C) identify peaks with a full-width-at-half-maximum of approximately 4-5 pixels (0.7-0.9 µm), the known diameter of oral Streptococcus cells (Baron, Samuel and Patterson, Maria Jevitz, 1996).

### 6 Supplementary Movie: SL-RPU Unmixed Dental Plaque Smear

3-D volume-rendered movie of a dental plaque smear labeled with 8 taxon-specific FISH probes (See Figure 6 in Main Text for color legend). Spectral image was unmixed with SL-RPU.
